# MetaR: simple, high-level languages for data analysis with the R ecosystem

**DOI:** 10.1101/030254

**Authors:** Fabien Campagne, William ER Digan, Manuele Simi

## Abstract

Data analysis tools have become essential to the study of biology. Here, we applied language workbench technology (LWT) to create data analysis languages tailored for biologists with a diverse range of experience: from beginners with no programming experience to expert bioinformaticians and statisticians. A key novelty of our approach is its ability to blend user interface with scripting in a single platform. This feature helps beginners and experts alike analyze data more productively. This new approach has several advantages over state of the art approaches currently popular for data analysis: experts can design simplified data analysis languages that require no programming experience, and behave like graphical user interfaces, yet have the advantages of scripting. We report on such a simple language, called MetaR, which we have used to teach complete beginners how to call differentially expressed genes and build heatmaps. We found that beginners can complete this task in less than 2 hours with MetaR, when more traditional teaching with R and its packages would require several training sessions (6-24hrs). Furthermore, MetaR seamlessly integrates with docker to enable reproducibility of analyses and simplified R package installations during training sessions. We used the same approach to develop the first composable R language. A composable language is a language that can be extended with micro-languages. We illustrate this capability with a Biomart micro-language designed to compose with R and help R programmers query Biomart interactively to assemble specific queries to retrieve data, (The same micro-language also composes with MetaR to help beginners query Biomart.) Our teaching experience suggests that language design with LWT can be a compelling approach for developing intelligent data analysis tools and can accelerate training for common data analysis task. LWT offers an interactive environment with the potential to promote exchanges between beginner and expert data analysts.

## INTRODUCTION

Present day biology often requires that biologists rely on software tools for data analysis. For instance, software tools are required for analysis of high-throughput data, for the study of genome-wide gene expression, genetic or epigenetic. Similarly, most fields of biology require specialized software tools for analysis of microscopy, crystallography or other data. Most analysis software is constructed in a very similar manner: writing a program as a collection of text source code compiled into one or more executable analysis tools. Despite the evolution of programming languages, encoding programs as text has been a constant since the invention of the first high-level programming language (FORTRAN Backus [1958, 1978]).

In this manuscript, we discuss several drawbacks of encoding programs as text that we believe make teaching of data analysis more difficult than necessary and contribute to some frequent challenges encountered by data analysts. Language Workbenches (LWs) with projectional editors offer an alternative platform to develop data analysis tools. LWs were conceived in the 90s Simonyi [1995] and have since led to the development of robust software development environments Dmitriev [2004], Erdweg et al. [2013]. In this study, we discuss an application of a LW platform to facilitate the teaching of data analysis to biologists. To this end, we used the Meta-Programming System (MPS, http://jetbrains.com/mps), a robust and open-source LW.

One question we were particularly interested in answering was whether we could create an analysis tool that blends the boundary between programming/scripting languages and graphical user interfaces, and therefore facilitate teaching complete beginners. Programming languages such as the R language Ihaka and Gentleman [1996] are frequently preferred for data analysis by experts. They have so far been the most flexible and powerful tools for data analysis, but require a steep learning curve. In contrast, beginners tend to prefer data analysis software with a graphical user interface, which are easier to learn, but eventually are found to lack flexibility and extensibility. We reasoned that blending these two types of interfaces into one tool could provide a simpler way for beginners to learn elements of scripting, improve repeatability and reproducibility of their analyses, and increase their productivity.

We found that LWT made it possible to quickly develop such a tool. We called this tool MetaR because it leverages the R ecosystem, but supports meta-programming. We designed this novel analysis tool using an iterative process that benefited from frequent feedback from users of the tool. In this manuscript, we describe the goals of the MetaR languages, explain how the tool can be used, and highlight the most innovative aspects of the languages compared to other tools used for data analysis, such as the R language Ihaka and Gentleman [1996] or electronic notebooks.

The initial focus of MetaR was on analysis of RNA-Seq data and the creation of heatmaps, but the tool is general and can be readily extended to support a broad range of data analyses. For instance, we have used MetaR to analyze data in a study of association between the allogenomics score and kidney graft function Mesnard et al. [2015]. We chose to focus on the construction of heatmaps as a use case and illustration for this study because this activity is of interest to many biologists who obtain high-throughput data.

We report on our experience teaching MetaR to complete beginners and compare the duration of such training sessions to that of similar training conducted with traditional approaches and tools. Importantly, we found that both beginners and experts can benefit from blending user interfaces and scripting. Beginners benefit because the MetaR user interface is much simpler to learn than the full R programming language. Expert users benefit because they can quickly prototype and develop high-level language elements to simplify repetitive aspects of data analysis.

## RESULTS

### Design of a High-level Data Analysis Language

Several decisions must be made when designing a new computational language. Most decisions are driven by design goals. We have designed the MetaR language to address the following goals:

1. The language should help users who have no knowledge of programming. The goal is to offer a smooth learning curve for beginners used to GUIs. We favor declarative language constructs over flexibility in parts of the language aimed at beginners.
2. Since a table of data is a frequent input when working with high-throughput data, make Table a first class element of the design. Leverage this element to simplify the annotation of the columns of a table. We rely on the idea that a little formalism (e.g., annotation of table columns) goes a long way to simplify analysis scripts.
3. Eliminate the need to know the language syntax to help beginners get started quickly. We leverage the MPS LW and its projectional editor to this end (Voelter and Solomatov [2010]). The MPS projectional editor provides interactive features, such as auto-completion, that provide guidance to beginners and experts alike when using the language to develop analyses.
4. Provide the ability to blend a scripting language with a graphical user interface. We use language composition and the ability of the MPS LW to render nodes with a mix of text and graphical user interface components.
5. Offer essential data transformations (e.g., joining two tables, taking subsets of rows of a table) via simple, yet composable language elements.
6. Provide means for experts to use their knowledge of the R language to work-around cases when the MetaR language is not sufficiently expressive to perform a specific analysis. We offer the ability to embed R code inside a MetaR analysis, as well as the ability to write scripts in the R language. In both instances, this variant of the R language supports language composition and enables embedding graphical user interfaces inside script fragments.

### Overview of MetaR and Composable R

Figure 1 presents an overview of the features offered by MetaR and Composable R, for the full range of users that the platform supports, from beginner to expert.

**Figure 1.**
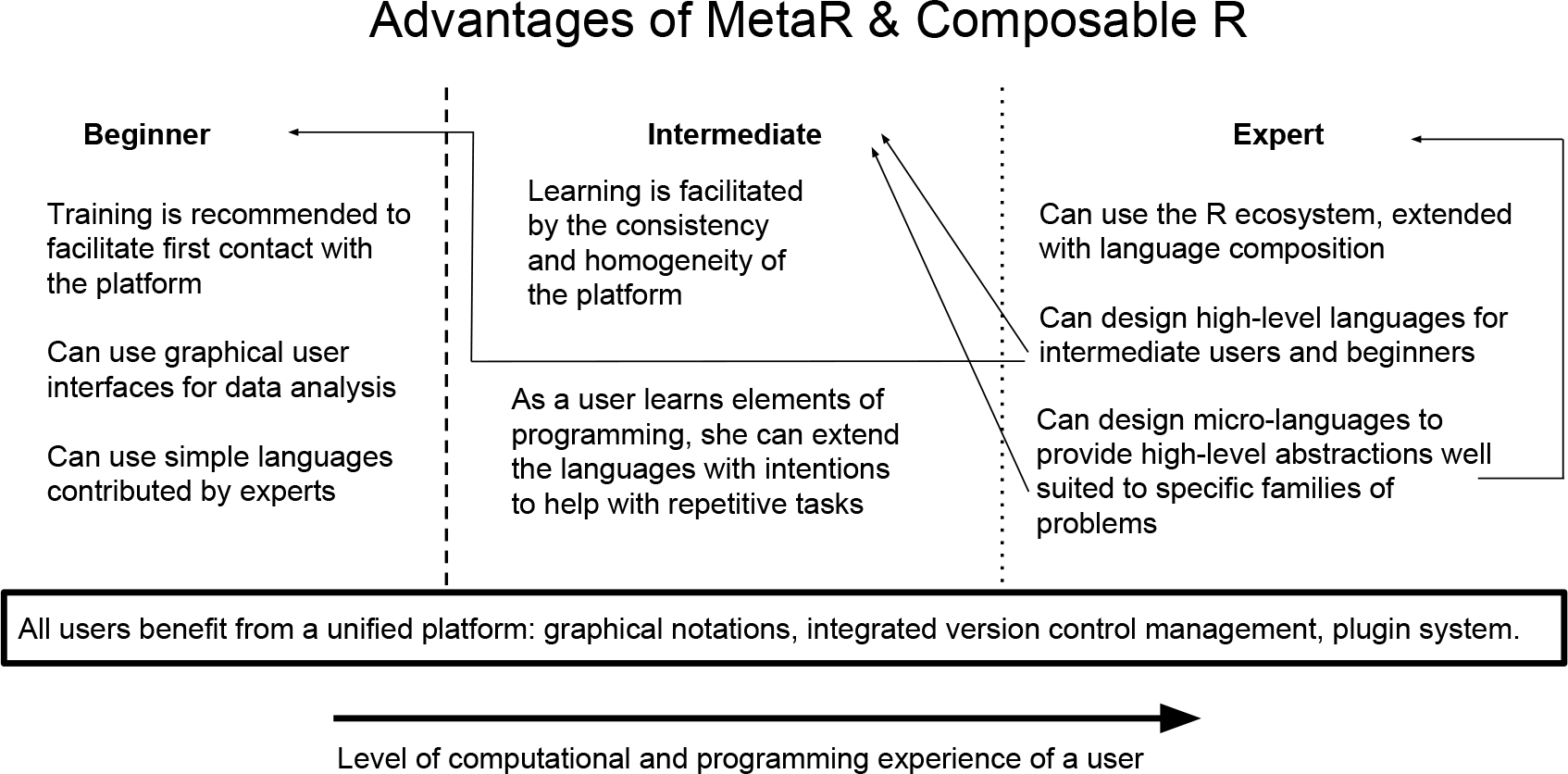
Overview of features provided by MetaR and Composable R in the LWT platform. We present the capabilities of the MetaR platform organized by the level of experience of a user. Beginners mostly benefit from the ability to blend graphical interfaces with scripting, and from high-level languages developed by experts on the platform. Intermediate users, who have basic programming skills, are able to customize the languages in simple ways, such as by creating intentions to help with repetitive steps of analyses. Intentions are context-dependent actions that can be added to a language at runtime (see Simi and Campagne [2014] for illustrations). Experts are users with strong programming skills who have become familiar with LWT. They can create micro-languages to extend Composable R, or design entirely new data analysis languages to help beginners with analysis for new domains. Users at all levels benefit from LWT platform features, including seamless integration of the languages with version control (see Benson and Campagne [2015] for a discussion of the integration with version control).

### Teaching the MetaR data analysis language

Teaching a data analysis tool can smooth the learning curve and prevent un-necessary frustration that students could experience if they tried working with the tool on their own. Since MetaR is a new platform for data analysis, training is also important to help users get started with the software.

For this reason, we developed detailed training material for the MetaR language and have offered training sessions at the Weill Cornell Medical College since January 2015. These monthly training sessions were offered to technicians, students, post-doctoral fellows and faculty across the institutions of our Clinical Translational Science Center (including three research institutions and one undergraduate city college), but included participants from other institutions in NYC.

When advertising the training sessions, we explicitly indicated that participants required minimal prior computer experience (“Analyses are executed in the R language, but no knowledge of R is needed to use MetaR”, “No programming or UNIX skills are required.”). This drew a large participation from attendants who had never used the R language or the command line.

Despite their limited computational experience, most participants were able to follow the 2h training session^1^ and construct a heatmap on their laptop by the end of the training. Participants who could not complete the assignment were either unable to install the software on an outdated laptop operating system (for instance, some problems included outdated security that prevented connection to the Wifi network at our institution), or encountered installation problems for R packages that would not install on their laptops on the day of training. As we recognized these problems, we developed simpler ways to install software dependencies on user laptops and encouraged users to download the required software before the session. We have so far trained approximately 150 participants in using MetaR to call RNA-Seq differential expression and create a heatmap.

The duration of MetaR training (2h) compares favorably with training sessions offered with R and bioConductor packages. An R/bioConductor workshop offered at the Weill Cornell Medical College requires about 6 hours of training (three two hour sessions) in order to help beginners conduct similar analyses to those performed in the MetaR training session. We requested help from the community to try and to quantify the amount of time typical R/bioConductor training sessions require. We created an online survey that trainers and trainees could fill out anonymously (http://goo.gl/forms/3ZWESUgtmd). The response rate of this survey was low, but indicated that between 6 and 24 hours was considered by most teachers a typical amount of time needed to teach how to call differentially expressed genes and creating a heatmap with R/bioConductor (two answers listed 6 hours, one answer 8 hours, two listed 24 hours). Interestingly, one trainee who answered the survey noted that 40 hours were required to learn the same skills, suggesting that trainers may under-estimate the amount of time needed when complete beginners try to learn these skills with R/bioConductor. The responses to this community survey indicate that traditional approaches require several 2 hr training sessions for a total of 6-24 hr training.

Assuming the responses to this survey are representative, MetaR training session are 3 to 6 times shorter than traditional training sessions. These data strongly support the notion that simple languages like MetaR can facilitate the teaching of data analysis for specialized analysis tasks. In the reminder of the Results Section, we explain the design of MetaR in more detail and present the features of the platform useful to experts.

### High-level Design Choices

In addition to the design goals presented previously, the design of MetaR included several strategic choices. We now present these choices and their rationales:

**Choice of a Target Language and Runtime System** A language needs a runtime system to execute the code of programs written in the language. A possible choice for a runtime is to target another high-level language (such as Java, or C) but this would require implementing all aspects of data manipulation in the target language. Since the R language (Ihaka and Gentleman [1996]) is widely used for data analysis in biology, we considered using it as a runtime system. Experts biostatisticians and bioinformaticians have developed many R packages that implement advanced analysis for biological high-throughput data. These packages can be used to simplify the implementation of a runtime system for a new data analysis language. We therefore decided that the MetaR language would generate R code in order to take advantage of the packages developed in this language. This decision greatly simplified the implementation of the MetaR language because it removed the need to develop a custom language runtime system.

**Data Object Surrogates** MetaR makes extensive use of Data Object Surrogates (DOS, our terminology). A data object surrogate is an object that represents other data (the source data). The surrogate often contains only limited information from the original data source. The DOS contains just enough to facilitate referring to the source data in another context for the purpose of data analysis, but not as much as to represent the entire content of the data source in memory. A good example of DOS is the Table object, which stores information about the columns of a data file. The Table DOS describes the columns of the table, but does not store the data contained in the table. A DOS typically has a name which can be used to refer to the DOS and its source data inside a MetaR model. References to table DOS help users refer to the table as they develop an analysis. Our use of the MPS LW facilitates the creation of DOS. In the MPS LW, we model DOS as concepts of the language. For instance, the Table DOS is represented by a Table concept, whose instances can be created in a model as root nodes. DOS are also used in MetaR to represent plots.

**Immutable Data Objects** Many programming languages (of which C, C++, Java, Perl, Python and R are members) make it possible to define variables or objects whose values can be changed (so called mutable variables). While this provides flexibility, it is a frequent source of confusion for beginners until they have developed their own mental model of how program steps modify variable values. During the design of MetaR, we chose to offer immutable objects rather than mutable variables when possible. This makes MetaR analyses easier to reason about because the value of objects cannot be changed after the object is created. This design decision does not prevent adding mutable variables to the MetaR language, but simplifies initial learning of the language by complete beginners.

### Organization into Languages

We designed MetaR as a collection of MPS languages. The main language, org.campagnelab.metar is aimed at beginners with limited computational experience.

In the next section, we explain how the MetaR language can be used from the point of view of an end-user. This section also includes highlights of features that differ from the state of the art in data analysis. Please note that exhaustive reference documentation is available elsewhere (see Campagne and Simi [2015]) and the goal of the following paragraphs is to provide a sufficient introduction to data analysis with MetaR that readers can understand the impact of the innovations we tested in developing this tool.

### The MetaR Language

#### Tables

An example of an immutable DOS is the MetaR Table object. In MetaR, objects of type Table represent tabular data with columns and rows of data. An example of a MetaR Table is shown in Figure 2. A MetaR Table is associated to a data file that contains the actual data of the table in a Tab-Separated Value (TSV). The location of the data file can be specified using Variables (i.e., $`{project}`), which offer independence from the local file system structure, and are particularly useful when keeping analyses under source control). A table has columns. Columns have names and types, which determine how data in each column is used. Types of data include string, numeric, boolean and enumeration (a small number of pre-defined categories, such as Male and Female). Figure 2 presents a table of RNA-Seq read counts which was obtained from the Gene Expression Omnibus Seguin-Estevez et al. [2014] and annotated to enable analysis with MetaR.

**Figure 2.**
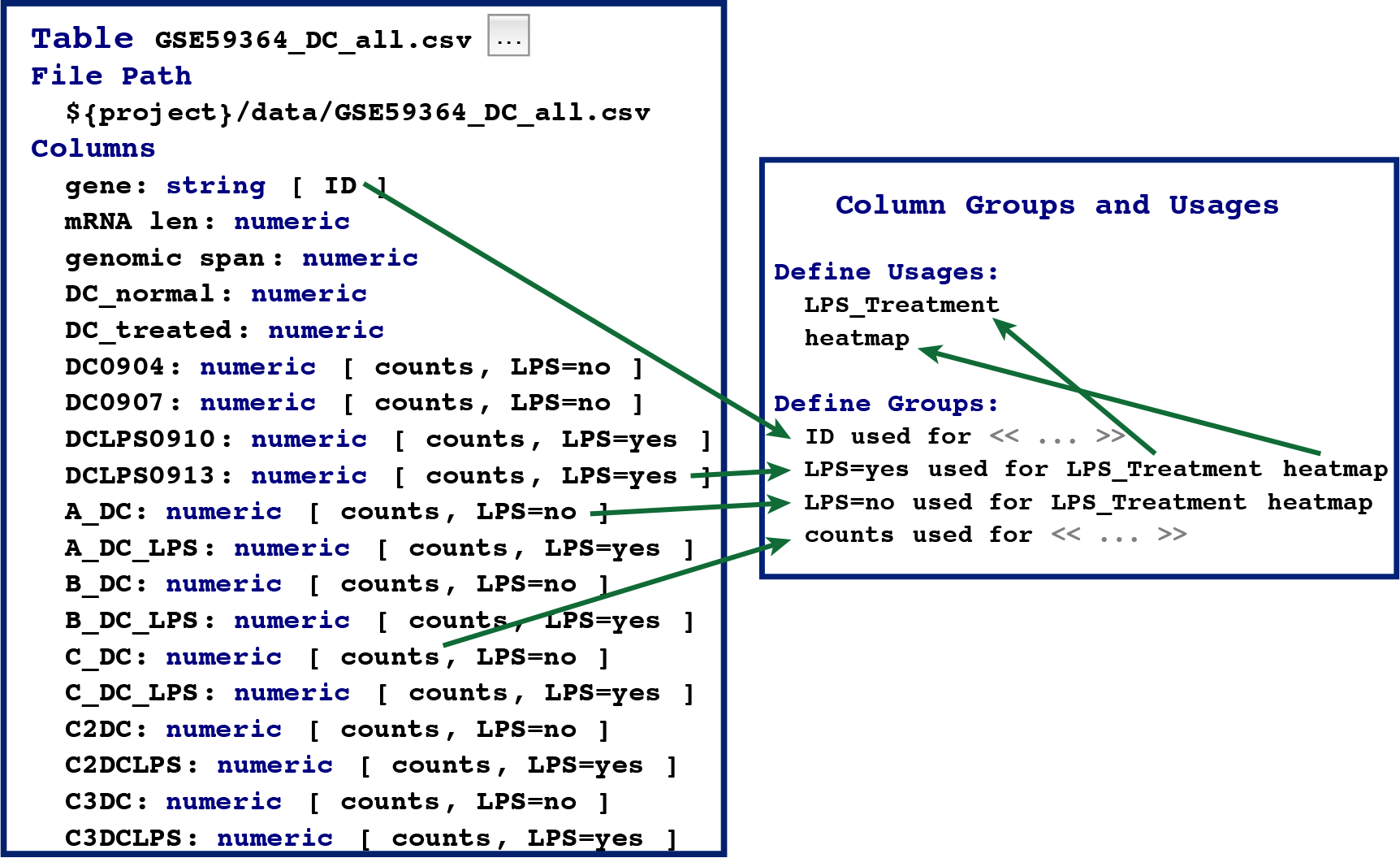
Table and Column Group objects. This figure presents the Table and Column Group objects. Green arrows show some cross-references among nodes of Tables and Column Groups. For instance, the ID group used to annotate the gene column is a reference to the ID group defined under the Column Group and Usage Container.

Annotating a Table consists of two steps: (1) browsing to the file that contains the data. This can be accomplished by clicking on the file dialog button (the little square with…) to locate the file. Upon selection of a valid file, the MetaR table node inspects the file and determines column names and types. Names and types are then shown in the Table node (under the `Columns` heading). (2) Specific columns can be annotated with one or more Column Groups.

Users can define arbitrary Column Groups in a different node called “Column Groups and Usages” (shown on the right of Figure 2). If two columns are related, user can define a Group Usage to explicitly document the relation. For instance, in Figure 2, the usage `LPS_Treatment` is defined to indicate that the Column Groups `LPS=no` and `LPS=yes` are two kinds of LPS treatments.

Tables and their annotations help users formalize information about data in a table. We find that asking the user to provide such information early on is beneficial because the structure of annotations can be leveraged in other parts of the language to provide intelligent auto-completion, customized for each table of data (for instance, to provide auto-completion for column names when writing expressions, or to select columns to use when joining two tables, examples of intelligent auto-completion is provided in the following sections, see Figure 3).

**Figure 3.**
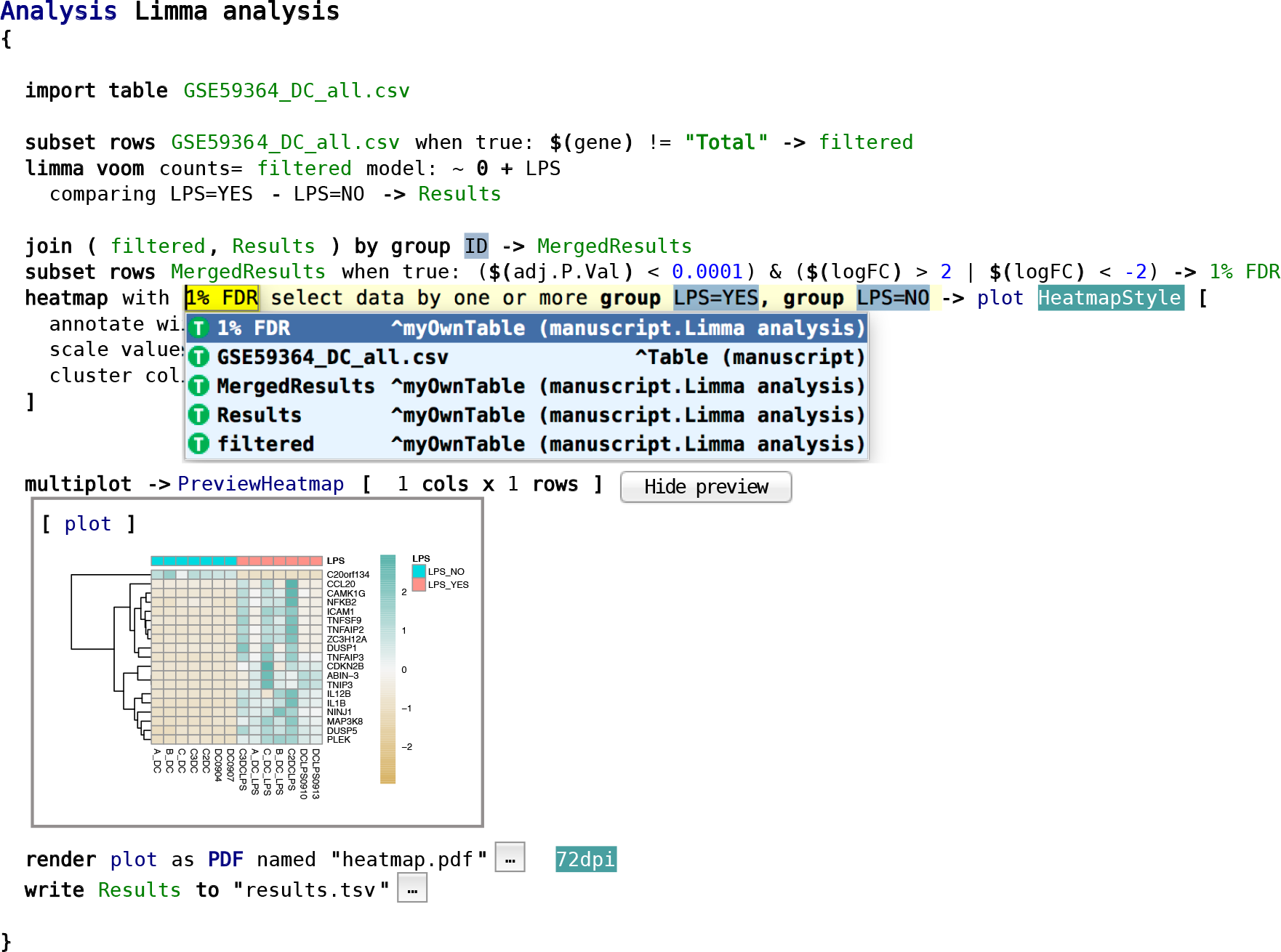
MetaR Analysis. The Analysis node is composed of a list of statements. This analysis works with the table of data presented in Figure 2, removes the row of data where the value `Total` appears in the gene column, performs statistical modeling with Limma Voom to identify genes differentially expressed between LPS treated and control samples, constructs a heatmap and displays the plot as a preview. Finally, the analysis converts the plot to PDF format and writes the joined table (statistics and counts) in the `results.tsv` file.

For instance, in the dataset of Seguin-Estevez et al. [2014], users can indicate which columns contain data for samples that were treated (`LPS=yes`) with lipopolysaccharide (LPS) or not (`LPS=no`). MetaR facilitates the data curation steps of a data analysis project by offering an interactive user interface to help users keep track of annotations. The interface is interactive in several ways: group names can be auto-completed to the groups defined in the “Column Group and Usage” object. Menus are available to add column group annotations to a set of columns that the user has selected. In addition to LPS treatment, Figure 2 shows the `count` annotation, used in an RNA-Seq differential expression analysis to identify which columns contain read counts, the ID column group, which uniquely identifies specific rows of the data table and the `heatmap` column group, used to choose which columns groups should be heatmap. This illustrates that the table annotation mechanism is flexible and can be leveraged by specific statements of the language, in order to indicate that the statement needs data annotated in a certain way.

#### Analyses

Analyses make it possible for users to express how data is to be analyzed. Figure 3 presents a MetaR Analysis node. This analysis is the one we use as a worked example during training sessions we offer at our institution. The editor of an analysis node offers an interface similar to that of a script in a traditional editor, but provides a more interactive and intelligent user interface. For instance, auto-completion is available at every point inside an analysis and suggests possible elements of the language that are compatible with the context at the cursor position.

The user may accept a suggestion and this results in the insertion of the language element at the position of the cursor. When the context calls for referencing a column of a table, for instance, only columns of Tables available at this point of the analysis are shown. While it is still possible to make mistakes when using this interface, mistakes created as a result of typos are less common than in programs encoded as text, for two reasons:

- Auto-completion offers a convenient way to set references between objects. Accepting an auto-completion suggestion helps users avoid typos.
- Some users choose not to use auto-completion to set references and instead type a referenced node name. In this case, mis-typed names that cannot be resolved to a valid node are highlighted in red and in the right margin of the editor (this feature of the MPS LW is available for all languages developed with the MPS platform). This highlighting draws the attention of the user to the error or typo. This feature is also important when merging different versions of an analysis placed under source control or when combining analyses from parts of other analyses (e.g., errors will be clearly marked after a code fragment is pasted into a new analysis).

Auto-completion help is available for the various types of references supported by the MetaR language. Examples of these can be seen on Figure 3 for tables (whose names are in green), plots (whose names are in blue), styles (names shown with a green background and white foreground, such as HeatmapStyle), or Column Group names (shown with a blue grey background and black foreground). Pressing control-B (or command-B on Mac) with the cursor on these nodes navigates to the destination of the reference (a menu is also available to help novice users discover this navigation mechanism). References may point to children nodes defined inside an analysis (e.g., plots), or nodes defined outside the analysis (e.g., tables and column groups).

Importantly, the MetaR user interface can also display buttons and images directly as part of the language. This feature takes advantage of the ability of the MPS LW to embed arbitrary graphical elements in the projectional editor. This capability is illustrated in Figure 3 by the “Hide Preview” button and by the heatmap image shown immediately below the `multiplot` keyword (pressing this button hides the plot preview).

The level of interactivity provided by the MetaR user interface is best conveyed by watching video recordings of its use. We provide training videos at http://metaR.campagnelab.org to illustrate how much more interactive the MetaR language is compared to other languages commonly used for data analysis.

### Language Composition and Micro-Languages

Since MetaR is implemented as a set of MPS languages, it fully supports language composition (Voelter and Solomatov [2010]). Language composition has no equivalent in text-based programming languages and many readers may be therefore unfamiliar with this technique. We will use an example to explain the advantage of this technique for data analysis.

Consider the table of results produced by the analysis shown in Figure 3. Users are likely to need to annotate the subset of genes found differentially expressed with gene names and gene descriptions. Information such as this is available in the Biomart system Haider et al. [2009].

To illustrate language composition, we created a new kind of MetaR statement called `query biomart`, which we defined in a micro-language. A micro-language is a language which provides only a few concepts meant to extend a host language. In this case, the MetaR language is the host language and `query biomart` is a concept contributed by the the micro-language. The purpose of this concept is to connect to Biomart and retrieve data. In the R language, this functionality is provided as a BioConductor package (called “biomaRt”, Durinck et al. [2005])

Querying Biomart in R consists in calling one of the functions defined in the package with specific parameters. The statement is very specialized, and for this reason would not typically be part of the core statements of a text-based programming language. Leveraging language composition, we can offer a dedicated statement that supports auto-completion in a remote Biomart instance. The statement acts as a specialized user interface designed to help users retrieve data from Biomart (in very much the same way that the web-based interface to Biomart helps users query this resource, but here completely integrated with the MetaR host language).

Figure 4 illustrates how the `query biomart` statement can be used to obtain gene annotations. In order to use these statements, end-users of MetaR would declare using both the *org.campagnelab.metar.tables* (the host language) and *org.campagnelab.metar.biomart* (the micro-language). In this specific case, the micro-language is provided with the MetaR distribution, but end-users can also implement other micro-languages to seamlessly combine them with the host language (the process for doing so is described in the MetaR documentation booklet Campagne and Simi [2015], Chapter 10). This capability makes it possible to customize the data analysis process for specific problems in much more flexible ways than would be possible with text-based programming languages: with the `query biomart` statement, we demonstrated that it is possible to remotely query databases to support auto-completion directly in the language. In contrast, text-based languages can only be extended in ways compatible with the syntax of the programming language, and are not able to support such levels of interactivity.

**Figure 4.**
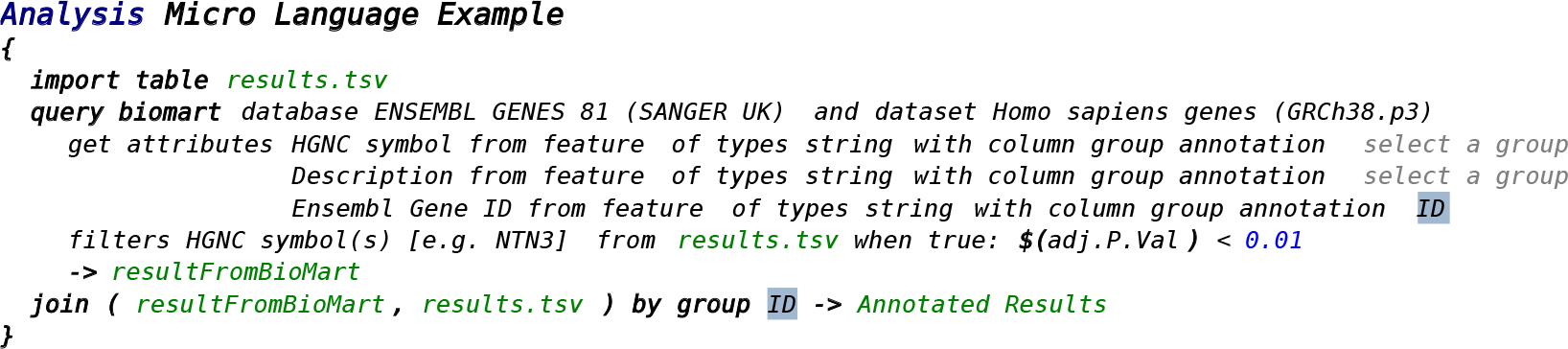
Example of Micro-language Composition. The query biomart stament is defined in a micro-language called *org.campagnelab.metar.biomart*, which extends the host language *org.campagnelab.metar.tables*. The biomart language provides one statement that offers an interactive user interface to help users retrieve data from biomart. This language reuses expressions and tables from the host language. Micro-languages can be enabled or disabled dynamically by the end-user at the level of a model. This example retrieves Human ENSEMBL identifiers and gene descriptions using the HGNC gene symbols used as identifiers in the Results table (see Figure 3 for the analysis that produced Results).

### Composable R language

In addition to the MetaR language illustrated in Figure 2-4, we have developed a composable R language. This language models the traditional R language Ihaka and Gentleman [1996], but supports language composition. Composable R is implemented in the language *org.campagnelab.metar.R* distributed with MetaR. R programs can be pasted in text form into an RScript root node and the text is parsed and converted to nodes of the composable R language. In Figure 5, we show the R code equivalent to the analysis shown in Figure 4. This R script was pasted from the text generated automatically from the MetaR analysis shown in Figure 4. Executing this script is supported in the MPS LW and yields the same result that of the simpler MetaR script shown in Figure 4.

**Figure 5.**
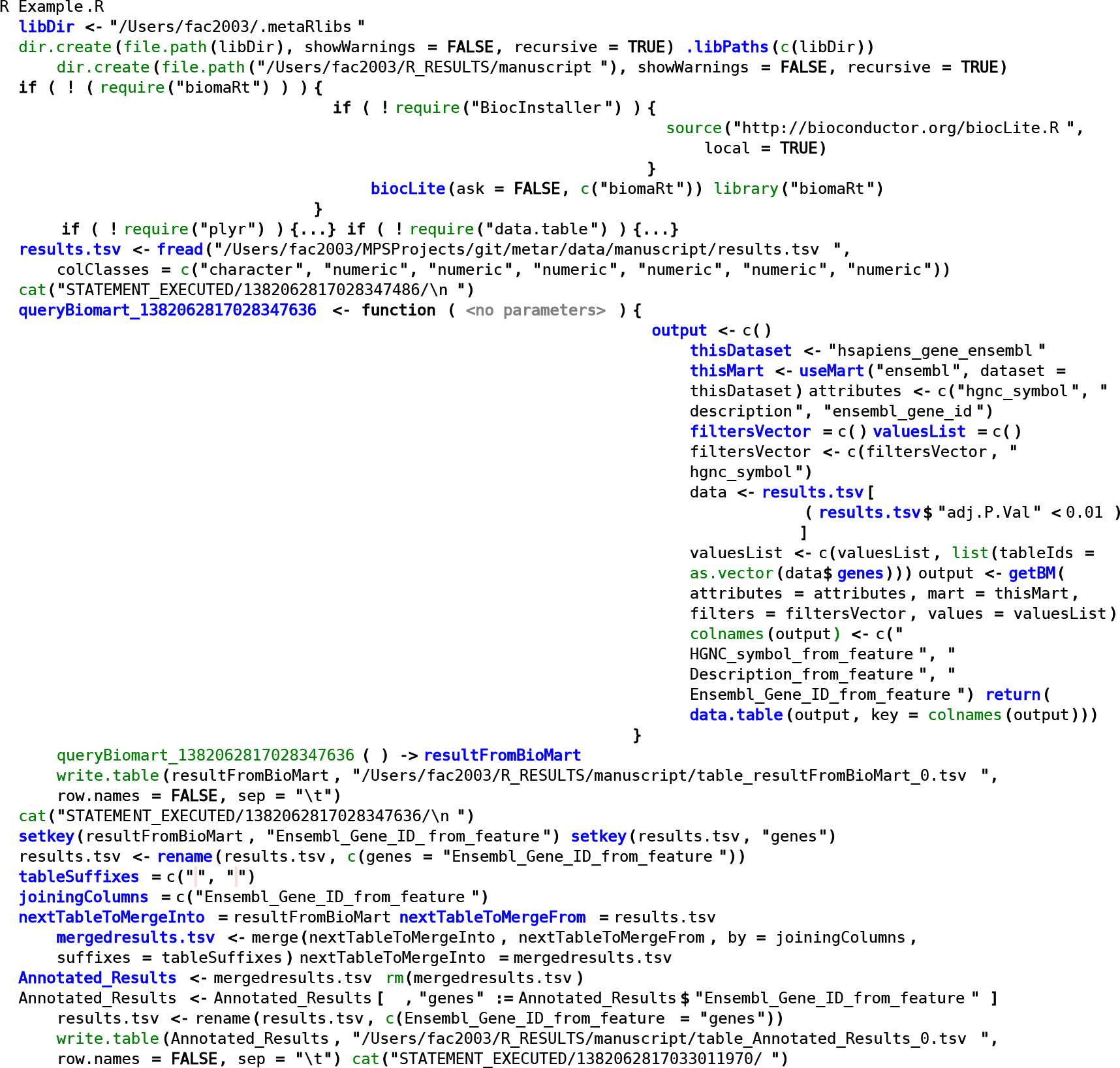
R language equivalent of the Analysis shown in Figure 4. To produce this figure, the analysis shown in Figure 4 was generated to the R language and the text was pasted in a RScript node of the composable R language. Automatic parsing of the R code into composable R objects yields a composable R version of the biomart example. Notice that boiler plate code needed to import R packages is shown only for the biomaRt package. Subsequent package import statements have been folded {…} to save space in the Figure. Folding is directly supported by the MPS LW. Function calls are highlighted in green and are linked to the function declaration in the package stub (end-user can navigate to each function to review its list of arguments, for instance). While it is likely that expert R programmers could produce somewhat more compact R code than this automatically generated code, comparison with Figure 4 indicates that a micro-languages can offer a concise alternative to a series of function call. The figure also illustrates the breadth of support for the language implemented in Composable R.

### Micro-Languages for the R Language

A composable R language makes it possible to create micro-languages that compose directly with R as the host language. We demonstrate this capability by adapting the `query biomart` statement shown in Figure 4 to the R language. Adaptation is simple because both MetaR and R generate to the same target language (R). In this case, we create a sub-concept of `Expr` (this type represents any R expression), and define a field of type `Biomart` (the concept that implements `query biomart`). This simple adapter is sufficient to make it possible to use the `query biomart` user interface inside an R script and is defined in the language org.campagnelab.metar.biomartToR. The result of composing the adapter language with composable R is shown in Figure 6. We also provide a short video to illustrate the interactive capabilities of a micro-language combined with composable R (see https://youtu.be/ZwGj1RPOODQ).

**Figure 6.**
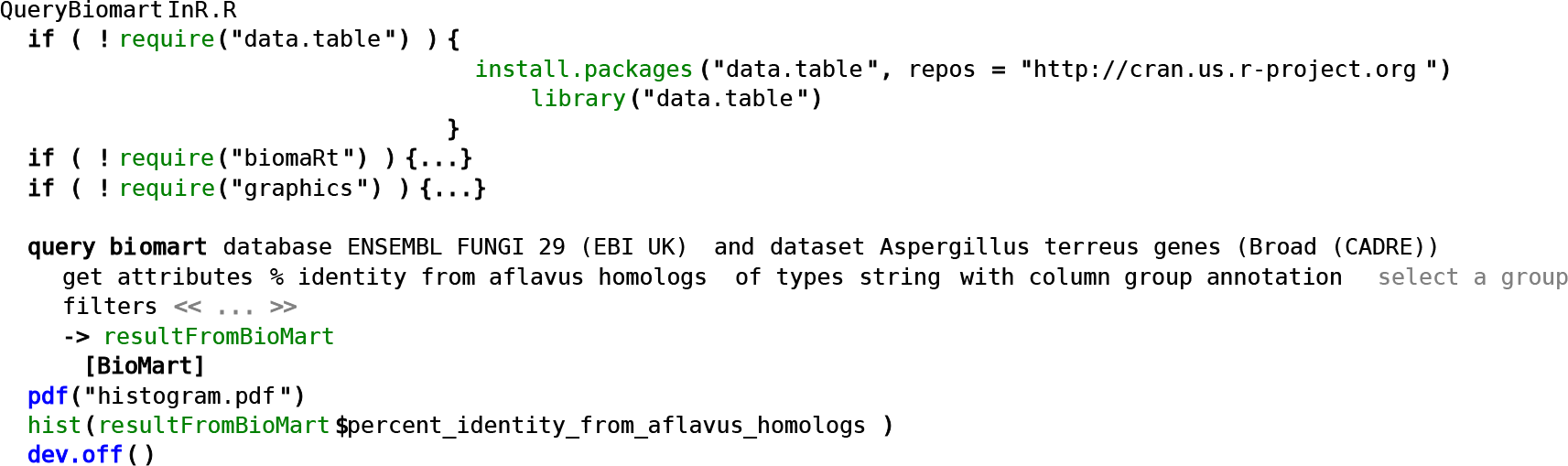
Composing Query Biomart with the composable R language. We developed an adapter that makes it possible to use the MetaR `query biomart` statement directly inside a composable R Script. This figure shows how the query biomart Expression adapter appears when used inside an R script. Notice how the table and column adapters are used inside a regular `hist()` function call `resultFromBioMart$percent_identity_from_aflavus_homologs`. These adapters make it possible to refer to the table produced by the statement as an R expression and provide auto-completion for column names in the table (determined dynamically based on the query expressed in the `query biomart` statement).

This example illustrates that a composable R language makes it possible to mix regular R code with new types of language constructs that can include user interfaces elements. This opens up new possibilities to facilitate repetitive analyses in R, for instance for specific data science domains (e.g., the Biomart example is useful for bioinformatic data analyses), but also for more general activities where simpler ways to perform a task would be beneficial. An example of this would be a micro-language to facilitate the use of packages to replace the boiler-plate package import code found at the beginning of most R scripts.

### Using R Expressions in the MetaR Language

Figure 7 illustrates that language composition can also be used to embed R expressions inside a MetaR analysis. This extension is possible because both analyses and R expressions generate code compatible with the syntax of the R programming language. Providing a way to embed the full language in a simpler analysis language offers a guarantee that the end-user will not be overly limited by restrictions of the simpler language.

**Figure 7.**
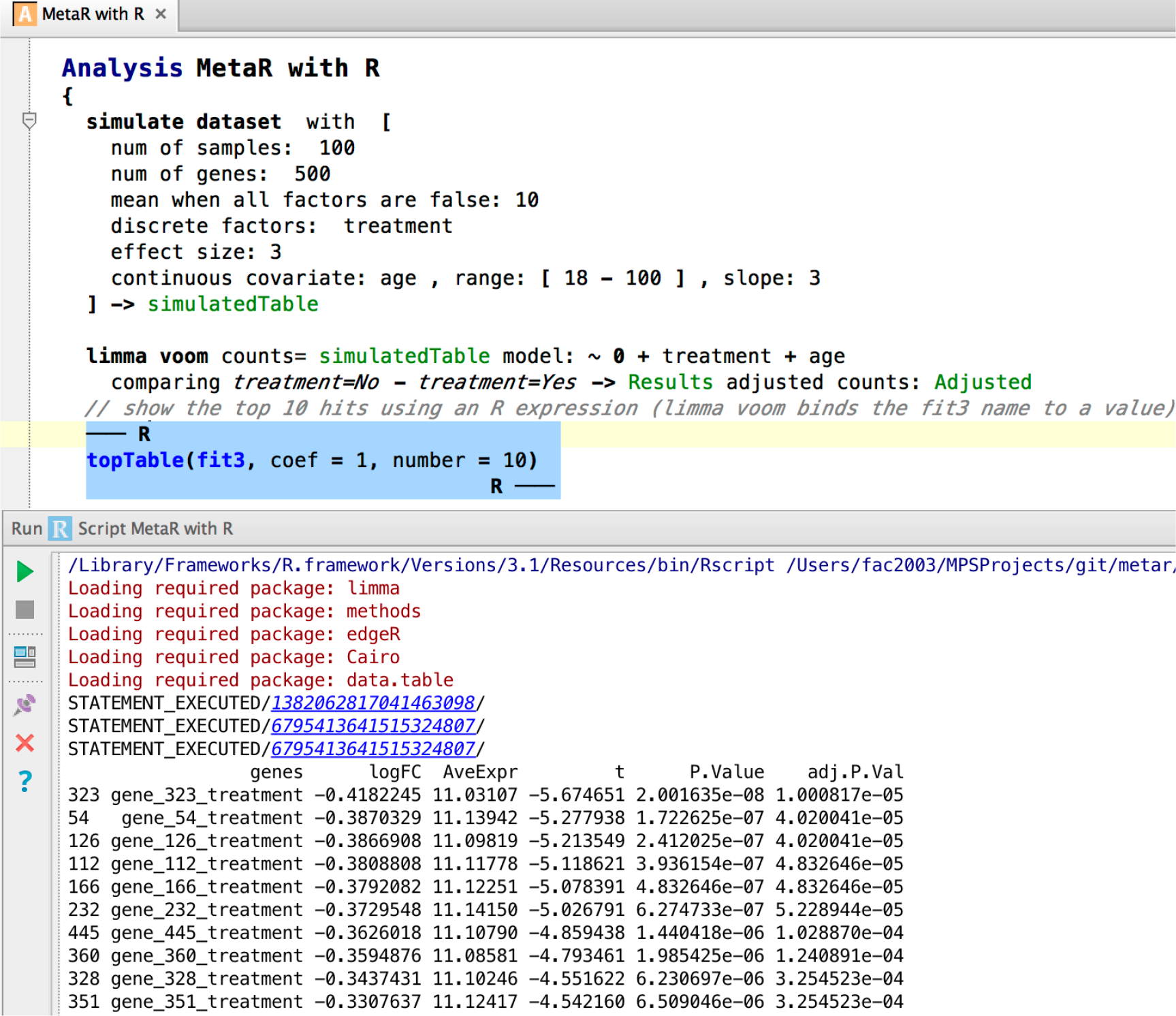
Composing R Expressions with the MetaR Language. Top panel: this example illustrates that it is possible to use R code inside a MetaR analysis. In this snapshot, R code is delimited by the — R and R — markers and shown with a blue background. Embedding R code in MetaR provides flexibility to perform operations for which MetaR statements have not yet been developed. The analysis shown simulates a dataset using simple parameters and tests the ability of Limma voom, as integrated with MetaR, to call differentially expressed genes. Bottom panel: shows the result of executing the analysis inside the MPS LW. As part of execution, the analysis is converted to R code, this code is run and standard output displayed inside the LW. The `STATEMENT EXECUTED//` lines hyperlink the progress of the execution with each specific analysis statement that has been executed.

### SOFTWARE

MetaR is distributed as a plugin of the MPS LW. Instructions for installing the software are available online at http://metaR.campagnelab.org. Briefly, after installing MPS, users can download and activate plugins with the Preferences/Plugins (Mac) or Settings/Plugins (Windows/Linux) menu. Plugins are stored as Zip files on the Jetbrains Plugin repository https://plugins.jetbrains.com/category/index?pr=mps&category_id=92 and can also be downloaded and installed manually from the zip file. Source code (technically, MPS languages serialized to files) are distributed on GitHub at https://github.com/campagnelaboratory/MetaR Campagne et al. [2015]. MetaR (and the MPS LW) are distributed under the open-source Apache 2.0 license.

## DISCUSSION

### Data Object Surrogates and Relation to Meta Data

DOS are related, but different from metadata. For instance, the Table DOS provides metadata about the file that contains the tabular data represented by Table nodes. It lists columns, associates columns to groups and defines group usages. This type of information can be thought of as metadata about the file that contains the tabular data. However, there is an important difference between DOS and metadata. For instance, a MetaR Table only provides metadata relevant to the analysis that the user needs to perform. It makes no effort to provide information that would have a meaning outside of the user’s analysis. This simplification maximizes the benefit of annotation while keeping the effort needed to produce it minimal and local to the user who actually needs the annotation.

### Graphical User Interfaces for Data Analysis

Programs with graphical user interfaces (GUIs) (also called direct manipulation interfaces Galitz [2007]) are often popular among beginners who are starting with data analysis and have no programming or scripting experience. GUIs are popular in part because they facilitate discovery of software functionality directly when using the software. They do not require prior-knowledge of syntax.

Data Analysis software with GUIs constrains how analysis is to be performed and provides clear menus and buttons that make it obvious what the program can do. A user can often discover new ways to perform analysis with these tools simply by browsing the user interface and looking at choices offered in menus and dialogs of the program. While such programs are favored by beginners (because they are relatively easy to learn), more advanced users who need to perform similar analyses across several datasets tend to strongly prefer analysis software that does not require repeating interactions with a GUI for every new dataset that must be studied. The novel approaches we have used to develop MetaR share these advantages with GUIs.

A minority of analysis software with GUIs also supports writing and running scripts in their user interface. For instance, JMP from SAS Inc. is an example of a statistical analysis software with GUI that also offers a scripting language. However, when scripting is offered, it is often only loosely integrated with the rest of the interface. Furthermore, users who are familiar with the GUI often need to learn scripting from scratch and do not benefit much from their prior experience using the GUI.

### Scripting and Programming Languages for Data Analysis

Scripting and programming languages are popular options for data analysis because analyses encoded in scripts or programs can be reused with different datasets. This makes these options popular among researchers who have programming skills and engage frequently in data analysis. The popularity and power of scripting for data analysis is epytomized by the development of the R language Ihaka and Gentleman [1996], which has become a defacto workhorse of open data science in biology. The versatility of the R language is its strength, but mastering the language requires elements of programming. Learning the R programming language is not as simple as learning how to use a GUI analysis tool and many users who would benefit from data analysis experience difficulties with the steep learning curve involved in learning programming and the R language.

In contrast to R, the MetaR language offers a much simpler alternative for users who have no prior programming background. At the same time, the Composable R language offers the means for expert R users to extend the R language with micro-languages in order to provide custom user interfaces. Such interfaces could be used to flatten the learning curve for novice data analysts or to empower expert data analysts with expressive means to encode solution to specific problems. Since both these options are available in the same platform (the MPS LW), users who become skilled with one language acquire transferable skills that help them learn other languages available on the platform.

### Impact on Development of User Proficiency

The MetaR high-level language shown in Figures 2 to Figure4 is aimed at novice data analysts. An interesting question is whether such a language can help novice data analysts learn skills that are useful when working with a variety of data analysis tasks.

If the language is sufficiently general, then novice users may learn skills that they can reuse when learning other general data analysis languages. If the language is too limited, then novice users would only learn a specialized analysis tool similar to existing GUI analysis tools. Rigorously determining to which category the MetaR language belongs would require following users for several months or years while they use the tool and we have not done such a study. However, we think that MetaR can help users transition to more general languages for the following reasons.

First, users who learn the high-level MetaR language acquire basic skills that are similar to those needed when working with other languages, including composable R. For instance, users learn to formalize their analysis intent using the constructs offered by the language. This is a very important first step that users with a strong programming background may take for granted, but that is difficult for novice users to acquire when they are distracted with problems of syntax. MetaR avoids syntax distractions and helps novice users focus on the logic of an analysis (e.g., how to combine language elements to achieve the desired analysis).

Second, the high-level MetaR language does not offer loops and conditionals. Since these language features are often needed for advanced analysis, many users who reach the point where they will need these language features will need to learn a language like R. MetaR offers composable R for this purpose. Novice users who have first learned the MetaR high-level language will be familiar with the MPS LW platform where composable R is also available. Some skills that users have acquired working with the high-level language will be directly transferable, including: how to run a script, how to navigate references to look at definitions, how to use auto-completion or use intentions to transform the program automatically, how to use source control (seamlessly integrated with the MPS LW). Subsets of the R language will still need to be learned to perform more advanced analysis, but learning can occur in an environment where the user is already comfortable. We believe that such an integrated environment where both high-level and low-level languages of the R ecosystem are offered will facilitate teaching of the many skills needed for data analysis. Formally testing whether this intuition is correct will require comparing cohorts of subjects learning data analysis. Alternatively, the answer may become apparent if a large number of data analysts were to transition to using composable R after initially learning the MetaR high-level language.

### Relation to Electronic Notebooks

MetaR shares some similarities to electronic notebooks such as IPython Pe´rez and Granger [2007], Jupyter (https://jupyter.org/) and Beaker (http://beakernotebook.com/) notebooks, but also has some important differences.

Regarding analogies, both MetaR and notebooks can be used to present analysis results alongside the code necessary to reproduce the results. For instance, the MetaR multi-view plot can be used to show a plot at the location where the statement is introduced in an analysis.

MetaR was developed approximately over the course of one year (2015). As such the software cannot be expected to be as feature-rich as software developed for many years. Beside this obvious difference, MetaR has the advantage to support language composition. In contrast, current data analysis notebooks support conventional programming languages constructed using text-based technology. Therefore, the closest that notebooks can approach language composition is to support multiple languages in one notebook, a so-called polyglot feature, available for instance in the Beaker notebook. Polyglot notebooks are useful, but cannot be extended by data analysts to customize languages for the requirements of a specific analysis project or domain. For instance, supporting a simple analysis language like MetaR would not be possible without developing a MetaR compiler and an associated execution kernel for the notebook. Developing and using micro-languages together with the traditional languages supported by the notebooks is also not possible.

Hence, the approach taken with MetaR is different from notebooks in two major ways. First, MetaR provides flexibility in designing new languages or micro-languages. It is not constrained by the syntax of a full programming language. Extending MetaR often consists in adding just one statement to an existing language. This promotes collaborative language design and development since many users can acquire sufficient skills to create one or two statements, reusing the building blocks provided by the host language (the steps needed to extend MetaR with a new language statement are described in the user manual Campagne and Simi [2015]). As long as a new statement generates valid R code, a MetaR Analysis that contains this statement will be executable.

Second, the syntax of the MetaR languages is not limited to text scripts or programs. Language Workbench technology used to implement MetaR supports graphical notations and diagrams as well as text. These differences combine to make it easier to design and implement custom data analysis abstractions with the LWT approach than it is possible with current electronic notebooks. Interestingly, the R IPython kernel could be used to execute scripts generated from MetaR analyses, which would provide an interactive console similar to that offered in the IPython notebook inside the MPS LW.

### Reproducible Research and Education

MetaR analysis and Composable R scripts can be executed seamlessly with an R environment installed inside a docker image (see Methods). Users can enable this feature by providing a few details about the installation of docker on their computer and checking the “Run with Docker” option in the MPS LW. This feature is particularly useful to facilitate reproducible research because docker images can be tagged with version numbers and always result in the same execution environment at the start of an analysis. This makes it possible to pre-install specific versions of R, CRAN and Bioconductor packages in a container and distribute this image with the MetaR analyses or R scripts that implement the analysis inside the container. While this is possible also with R, using docker on the command line, the customization of the MPS LW makes it seamless to run analyses with docker. We are not aware of a similar feature being supported by current R IDEs.

We found this feature also particularly useful for training sessions where installation of a working R environment can be challenging on trainees’ laptops. Using docker, we simply request that trainees pre-install Kitematic (available on Mac and Windows), or run docker natively on Linux and download the image we prepared with the packages used in the MetaR training sessions. The ability to run MetaR analysis in docker container results in a predictable installation of dependencies for training session and frees more of the instructor’s time to present data analysis techniques.

## METHODS

### Language Workbench Technology Primer

Since many readers may not be familiar with LWT, this section briefly describes how this technology differs from traditional text-based technology.

Text-based programming languages are implemented with compilers that internally convert the text representation of the source code into an abstract syntax tree (AST), a data structure used when analyzing and transforming programming languages into machine code.

In the MPS LW, the AST is also a central data structure, but the parsing elements of the compilers are replaced with a graphical user interface (called a projectional editor) that enables users to directly edit the data structure. Where text-based languages are restricted to programs written as text, a projectional editor can support both textual and graphical user interfaces (such as images, buttons, tables or diagrams) Voelter and Solomatov [2010]. Projectional editors can also offer distinct views of the same AST, implemented as alternative editors. Projectional editors keep an AST in memory until the user saves the program. Saving an AST to disk is done using serialization (loading is conversely done via deserialization to memory AST data structures).

The choice of serialization rather than encoding with text has a profound consequence. Serialization uniquely identifies the concept for each node in an AST. This method makes it possible to combine AST fragments expressed with different languages, when the concept hierarchy of the languages supports composition. We have presented examples of language composition in Simi and Campagne [2014], Benson and Campagne [2015]. In this manuscript, we extensively use language composition to extend the R language and provide the ability to embed user interfaces into R programs.

### Abstract Syntax Tree (AST)

An AST is a data structure traditionally used by compilers as a step towards generating machine code. In the MPS Language Workbench, an AST is a tree data structure, where nodes of the tree are instances of concepts (in the object-oriented sense). Figure 8 illustrates the notion of AST nodes, concepts and projectional editor.

**Figure 8.**
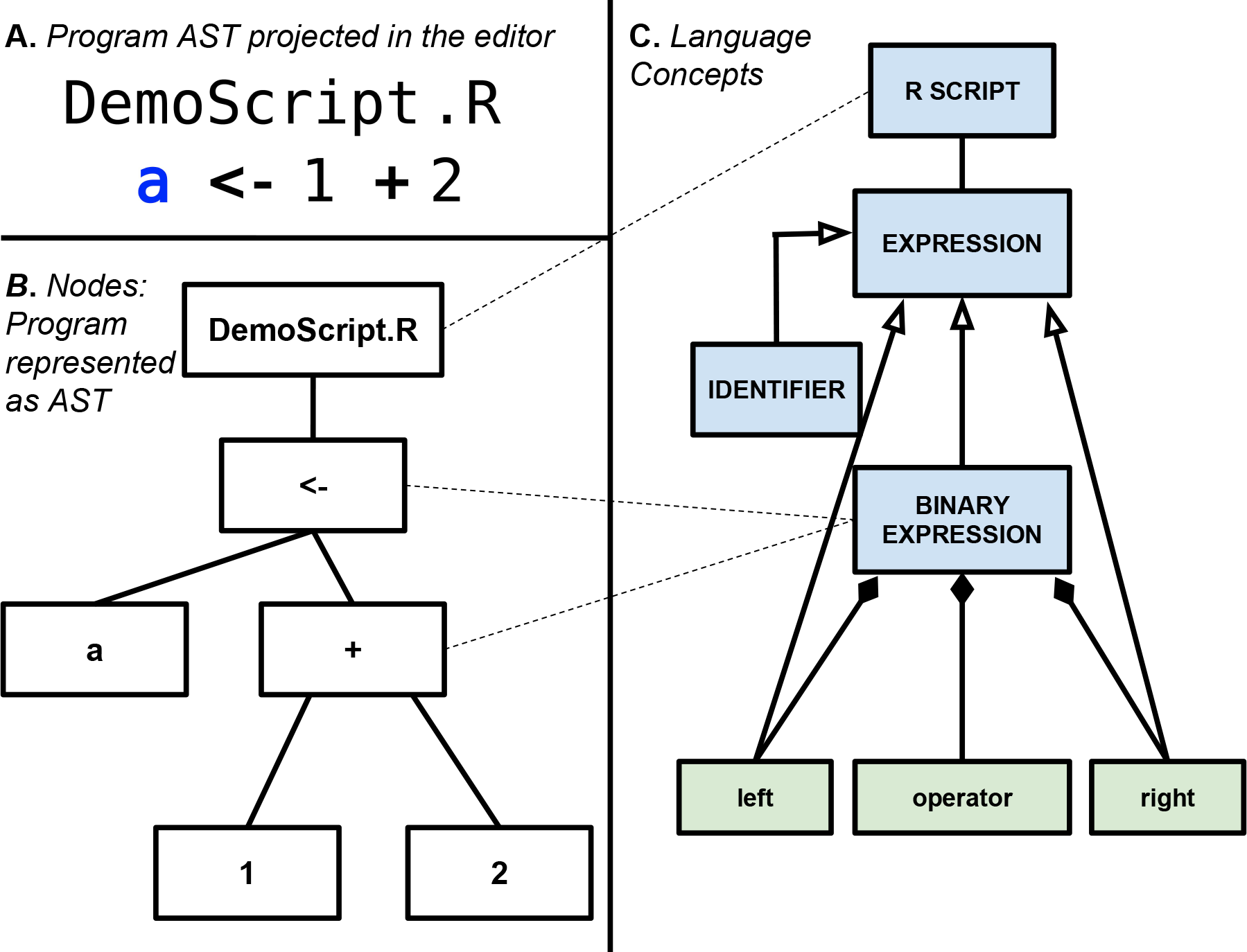
Concepts, Nodes and Projectional Editor. Panel A: Projectional editor showing a simple R script with one assignment expression. Panel B: An abstract syntax tree is shown with nodes that correspond to the program in panel A. Panel C: Language Concepts for the nodes in Panel (B) (shown as blue boxes). Each concept is connected to other concepts with an open-ended arrow to indicate inheritance (e.g., A <- B indicates that B is a sub-concept of A). Green boxes indicate fields of a concept and are connected to the concept that has these fields by a line with a black diamond on the concept that owns the field. This shows that `BinaryExpression` is a concept that is an `Expression` and has three fields: `left, operator` and `right`. Dotted lines connect nodes to their concept. For instance, the <- and + nodes are instances of `BinaryExpression`.

AST concepts may have properties (values of primitive types), children (lists of other nodes they contain), references (links to other nodes defined elsewhere in the AST). An AST has always a root node, which is used to start traversing the tree. In the MPS LW, AST root nodes are stored in models.

### Languages

In the MPS LW, languages are defined as collection of concepts, concept editors (which together implement the user interface for the language), and other language aspects Campagne [2014]. Each language has a name which is used to import, or activate, the language inside a model. After importing a language into a model, it becomes possible to create ASTs with this language in the model. Creating an AST starts with the creation of a root node. Children of the root node are added using the projectional editor. Children of root nodes, properties and references can be edited interactively in the editor.

We have used the MPS Language Workbench (http://jetbrains.com/mps), as also described in Campagne [2014] and Campagne [2015]. For an introduction to Language Workbench Technology (LWT) in the context of bioinformatics see Simi and Campagne [2014] and Benson and Campagne [2015] in the context of predictive biomarker model development.

### Language Design

We designed the MetaR MPS languages through an iterative process, releasing the languages at least weekly to end-users at the beginning of the project and adjusting designs and implementations according to user feedback. Full language developments logs are available on the GitHub code repository (https://github.com/CampagneLaboratory/MetaR) Campagne et al. [2015]. Briefly, we designed abstractions to facilitate specific analyses and implemented these abstractions with the structure, editor, constraints and typesystem aspects of MPS languages. Generated R code is produced from nodes of the languages using the org.campagnelab.TextOutput plugin. An illustration of the steps required to develop one language statement is available in Chapter 10 of the MetaR documentation booklet (see Campagne and Simi [2015]).

### Table Viewer

We implemented a Table viewer as an MPS Tabbed Tool, using the MPS LW mechanisms for user interface extension (see Campagne [2015]). The table viewer provides the ability to inspect the data content of any table produced during an analysis, or any input table. When the cursor is positioned over a node that represent a FutureTable (created when running the R script generated from the MetaR Analysis), and the viewer is opened, it tries to load the data file that the analysis would create for this table. If the file is found, the content is displayed using a Java Swing Component in the MPS user interface of the Table Viewer tool.

### Language Execution

MetaR analyses can be executed directly from within the MPS LW. This capability was implemented with Run Configurations (see Campagne [2015], Chapter 5).

### Execution in a Docker Container

In order to facilitate reproducible execution, we implemented optional execution within a Docker container. A docker image was created to contain a Linux operating system and a recent distribution of the R language (provided in the rocker-base image), as well as several R packages needed when executing the MetaR statements. The Run Configuration was modified to enable execution inside a docker container when the user selects a checkbox “execute inside docker container”. Information necessary to run with docker (i.e., location of the docker executable, docker server connection settings and image name and tag) is collected under a tab in the MPS Preferences (Other Settings/Docker).

## Footnotes

1 Sessions are scheduled for 2hrs, but often complete half an hour early when participants do not require software installation troubleshooting at the start of the session.

## Acknowledgments

We thank the authors of the `rocker-base` image, used in the MetaR project to facilitate the creation of docker images for training sessions and reproducible research. This investigation was supported by the National Institutes of Health NIAID award 5R01AI107762-02 to Fabien Campagne and by grant UL1 RR024996 (National Institutes of Health (NIH)/National Center for Research Resources) of the Clinical and Translation Science Center at Weill Cornell Medical College. Declaration of competing interests: Fabien Campagne is the author of the books “The MPS Language Workbench, Volume I and II” and receives royalties from the sale of these books.

